# Untangling featural and conceptual object representations

**DOI:** 10.1101/607499

**Authors:** Tijl Grootswagers, Amanda K. Robinson, Sophia M. Shatek, Thomas A. Carlson

## Abstract

How are visual inputs transformed into conceptual representations by the human visual system? The contents of human perception, such as objects presented on a visual display, can reliably be decoded from voxel activation patterns in fMRI, and in evoked sensor activations in MEG and EEG. A prevailing question is the extent to which brain activation associated with object categories is due to statistical regularities of visual features within object categories. Here, we assessed the contribution of mid-level features to conceptual category decoding using EEG and a novel fast periodic decoding paradigm. Our study used a stimulus set consisting of intact objects from the animate (e.g., fish) and inanimate categories (e.g., chair) and scrambled versions of the same objects that were unrecognizable and preserved their visual features (Long, Yu, & Konkle, 2018). By presenting the images at different periodic rates, we biased processing to different levels of the visual hierarchy. We found that scrambled objects and their intact counterparts elicited similar patterns of activation, which could be used to decode the conceptual category (animate or inanimate), even for the unrecognizable scrambled objects. Animacy decoding for the scrambled objects, however, was only possible at the slowest periodic presentation rate. Animacy decoding for intact objects was faster, more robust, and could be achieved at faster presentation rates. Our results confirm that the mid-level visual features preserved in the scrambled objects contribute to animacy decoding, but also demonstrate that the dynamics vary markedly for intact versus scrambled objects. Our findings suggest a complex interplay between visual feature coding and categorical representations that is mediated by the visual system’s capacity to use image features to resolve a recognisable object.

## Introduction

How does the brain transform perceptual information into meaningful concepts and categories? One key organisational principle of object representations in the human ventral temporal cortex is animacy (Caramazza & Mahon, 2003; Caramazza & Shelton, 1998; Kiani, Esteky, Mirpour, & Tanaka, 2007; Kriegeskorte et al., 2008; Mahon & Caramazza, 2011; Spelke, Phillips, & Woodward, 1995). Operationalised as objects that can move on their own volition, animate objects evoke different activation patterns than inanimate objects in human brain activity patterns in fMRI (Cichy, Pantazis, & Oliva, 2014; Connolly et al., 2012; Downing, Jiang, Shuman, & Kanwisher, 2001; Grootswagers, Cichy, & Carlson, 2018; Konkle & Caramazza, 2013; Kriegeskorte et al., 2008) and in MEG/EEG (Carlson, Tovar, Alink, & Kriegeskorte, 2013; Contini, Wardle, & Carlson, 2017; Grootswagers, Ritchie, Wardle, Heathcote, & Carlson, 2017; Grootswagers, Robinson, & Carlson, 2019; Kaneshiro, Guimaraes, Kim, Norcia, & Suppes, 2015; Ritchie, Tovar, & Carlson, 2015). A current theoretical debate concerns the degree to which categorical object representations in ventral temporal cortex are due to systematic featural differences within categories (Long et al., 2018; op de Beeck, Haushofer, & Kanwisher, 2008; Proklova, Kaiser, & Peelen, 2016).

Recent work has focused on understanding the contribution of visual features to the brain’s representation of categories, such as animacy. This work has shown that a substantial proportion of animacy (de)coding in ventral temporal cortex can be explained by low and mid-level visual features (e.g., texture and curvature) that are inherently associated with animate versus inanimate objects (Andrews, Watson, Rice, & Hartley, 2015; Bracci & Op de Beeck, 2016; Bracci, Ritchie, & de Beeck, 2017; Bracci, Ritchie, Kalfas, & Op de Beeck, 2019; Coggan, Liu, Baker, & Andrews, 2016; Kaiser, Azzalini, & Peelen, 2016; Long et al., 2018; Proklova et al., 2016; Rice, Watson, Hartley, & Andrews, 2014; Ritchie, Bracci, & op de Beeck, in press; Watson, Young, & Andrews, 2016). Long et al. (2018) recently investigated how mid-level features contribute to categorical representations using images of intact objects and scrambled “texform” versions of the same objects. Crucially, the texform versions of the objects were unrecognisable (at the individual image identity level) but preserved mid-level features such as texture. Using fMRI, they found the categories of animacy and size were similarly coded in the brain for intact and texform versions of objects, thus demonstrating that such patterns can arise without the explicit recognition of an object (Long et al., 2018). In MEG and EEG, one study showed that animate and inanimate objects cannot be differentiated when they are closely matched for shape (Proklova, Kaiser, & Peelen, 2019). Other studies, however, have found that object animacy decoding generalises to unseen exemplars with different shapes (cf. Contini et al., 2017), suggesting animacy decoding, in part, might be based on general conceptual representations. Taken together, these results suggest that either there is some abstract conceptual representation of animacy, or that objects within the animate and inanimate categories share sufficient visual regularities to drive the categorical organisation of object representations in the brain.

In the current study, we tested the contribution of visual features to the dynamics of emerging conceptual representations. We used a previously published stimulus set (Figure 1) that was designed to test the contribution of mid-level features to conceptual categories (animacy and size) in the visual system (Long et al., 2018), which consisted of luminance-matched real objects, and scrambled, “texform” versions of the same objects that retain mid-level texture and form information (Long, Störmer, & Alvarez, 2017; Long et al., 2018). We used EEG and a rapid-MVPA paradigm (Grootswagers et al., 2019) to study the emergence of conceptual information. Based on previous fMRI work (Long et al., 2018), we predicted that texforms would evoke animacy-like patterns in the EEG signal similar to intact objects. In addition, we hypothesized that animacy-like patterns evoked by texforms may need more time to develop. To test this, we presented the stimuli at varying rapid presentation rates, as faster rates have been shown to limit the depth of stimulus processing (Collins, Robinson, & Behrmann, 2018; Grootswagers et al., 2019; McKeeff, Remus, & Tong, 2007; Robinson, Grootswagers, & Carlson, 2019). We found that EEG activation patterns of texform versions of the objects were decodable, but that conceptual categorical decoding of intact objects was more robust, and could be achieved at faster presentation rates, which suggests that the visual system needs less time to process the intact objects. Together, our results provide evidence that visual features contribute to the representation of conceptual object categories, but also show that higher level abstractions cannot be fully explained by statistical regularities.

**Figure 1.**
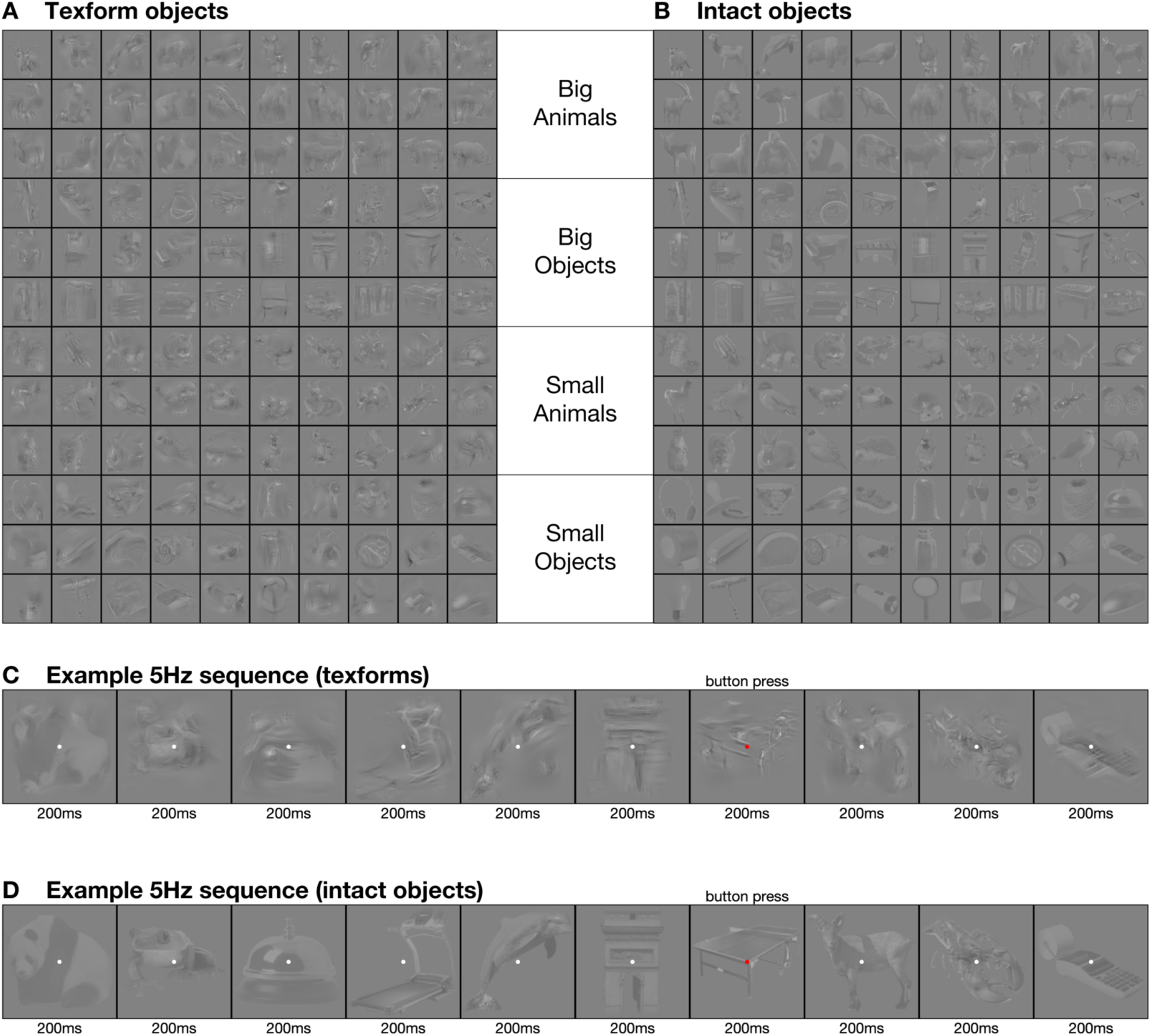
Stimuli and design. Stimuli were 120 objects categorizable as animate or inanimate, and as big or small. A. The first half of the experiment used texform versions of objects (presented first so that participants were not aware of their intact counterparts). B. In the second half of the experiment, the original intact versions were used. All images were obtained from https://osf.io/69pbd/ (Long et al., 2018). Stimuli were presented at four presentation frequencies. C. Example texform sequence at 5Hz, where stimuli were presented for 200ms each. D. Example intact object sequence at 5Hz. The sequence presentation orders for intact objects and texforms were matched. Participants performed an orthogonal task where they responded with a button press to the fixation dot turning red.

## Methods

Stimuli, data, and analysis code are available online through https://osf.io/sz9ve.

### Participants

Participants were 20 volunteers (11 females, 9 males; mean age 24.6, age range: 17-59) recruited from the University of Sydney in return for payment or course credit. All participants reported normal or corrected-to-normal vision. Two participants were left-handed. The study was approved by the University of Sydney ethics committee and informed consent in writing was obtained from all participants.

### Stimuli and design

Stimuli were obtained from https://osf.io/69pbd (Long et al., 2018). For a full description of the stimulus generation procedures, see Long et al., 2018. The stimuli were 120 visual objects that were grouped in four categories: big animals, small animals, big objects, and small objects. This allowed for orthogonal animacy and size categorisation of the stimuli. All stimuli were matched for average luminance. The stimuli underwent a scrambling procedure (Long et al., 2018) to generate texform versions of the same objects. All 240 stimuli, 120 intact objects, and 120 texform versions were used in this experiment (Figure 1).

Following the procedure of Long et al., we presented participants with texform versions of the stimuli in the first half of the experiment, and with intact objects in the second half of the experiment. Participants were all naïve to the experiment aims and were not informed about the relationship between the texforms and intact images. We used a rapid serial visual processing paradigm to present the stimuli in fast succession (Grootswagers et al., 2019). Stimuli were presented in random order in streams at four presentation frequencies: 60Hz, 30Hz, 20Hz, and 5Hz, always using a 100% duty cycle, following previous work that investigated category decoding at fast presentation rates (Grootswagers et al., 2019; Mohsenzadeh, Qin, Cichy, & Pantazis, 2018). All stimuli within a category (texforms/objects) were presented in each stream (i.e., every stream contained 120 images). Stimuli were presented at 6.8 x 6.8 degrees of visual angle on a grey background and were overlaid with a white fixation dot of 0.2 degrees diameter (Figure X). During the experiment, participants responded with a button press when the dot changed colour, which happened between 1 and 4 times during each stream, at random positions in the stream. Each object was presented 30 times in each condition (intact and texform), and at each presentation frequency. The experiment lasted about 40 minutes.

### EEG recordings and preprocessing

Continuous EEG data were recorded from 64 electrodes arranged according to the international standard 10–10 electrode placement system (Oostenveld & Praamstra, 2001) using a BrainVision ActiChamp system, digitized at a 1000-Hz sample rate and referenced online to Cz. Preprocessing was performed offline using EEGlab (Delorme & Makeig, 2004). Data were filtered using a Hamming windowed FIR filter with 0.1Hz highpass and 100Hz lowpass filters and were downsampled to 250Hz. No further preprocessing steps were applied. All analyses were performed on the channel voltages at each time point. Epochs were created for each stimulus presentation ranging from [-100 to 1000 ms] relative to stimulus onset.

### Decoding analysis

We applied an MVPA decoding pipeline (Grootswagers, Wardle, & Carlson, 2017) applied to the EEG channel voltages. The decoding analyses were implemented in CoSMoMVPA (Oosterhof, Connolly, & Haxby, 2016). Regularised linear discriminant analysis (LDA) classifiers were used in combination with a sequence cross-validation approach to decode pairwise image identities firstly between all pairs of texform images, and secondly between all pairs of intact object images. For animacy decoding, an exemplar-by-sequence cross-validation approach was used (Carlson et al., 2013; Grootswagers et al., 2019). That is, a pair of animate and inanimate images from one sequence was used as test data, and classifiers were trained on the remaining images from the remaining sequences. This was repeated for all animate-inanimate pairs and all sequences, averaging the resulting cross-validated prediction accuracies. Real-world size decoding used the same pipeline, with an exemplar-by-sequence cross-validation procedure. To test for the similarity between texform and object patterns, we performed the same analyses and cross-validation designs described above, but we trained the classifiers on intact object sequences and tested on the texform sequences, and vice versa. All analyses were repeated for each time point in the epochs, resulting in a decoding accuracy over time for every presentation frequency, within subject. The subject-averaged results for each frequency were analysed at the group level.

### Statistical inference

For each decoding analysis, we used Bayesian statistics to determine the evidence for above chance decoding or non-zero differences between texform and intact object decoding accuracies (Dienes, 2011, 2016; Jeffreys, 1961; Rouder, Speckman, Sun, Morey, & Iverson, 2009; Wagenmakers, 2007). For the alternative hypothesis of above-chance (50%) decoding or a non-zero difference, a JZS prior (Rouder et al., 2009) was set with a scale factor of 0.707 (Jeffreys, 1961; Rouder et al., 2009; Wetzels & Wagenmakers, 2012; Zellner & Siow, 1980). We then calculated the Bayes factor (BF) which is the probability of the data under the alternative hypothesis relative to the null hypothesis. We thresholded BF > 10 as evidence for the alternative hypothesis, and BF < 1/3 as evidence in favour of the null hypothesis (Jeffreys, 1961; Wetzels et al., 2011). In addition to the Bayes factors, we computed p-values for decoding against chance, and the differences between texform and intact object decoding accuracies. We used a sign-swap permutation test (1000 iterations), and computed threshold-free cluster enhancement (TFCE; Smith & Nichols, 2009) values at each time point. To correct for multiple comparisons, the maximum TFCE statistic across time for each permutation were selected to form a corrected null-distribution (Maris & Oostenveld, 2007). We then calculated p-values by comparing the observed TFCE values to the corrected permutation distribution.

### Exploratory channel-searchlight analysis

To obtain insights into the source of the difference between texform and intact object decoding, we performed a channel by timepoint searchlight. For all contrasts, we performed multiclass decoding instead of all pairwise comparisons to reduce computation time. A leave-one-sequence-out cross-validation approach was performed on local clusters of channels. For each channel, a local cluster was constructed by taking the closest four neighbouring channels, and the decoding analyses were performed on the signal of just these channels. The decoding accuracies were stored at the centre channel of the cluster. This resulted in a time by channel map of decoding accuracy for each of the contrasts, and for each subject, at each frequency. Here, we reported the results for the decoding differences at 5Hz and have added the other frequencies to the project’s online repository.

## Results

Participants (N=20) viewed streams of texform stimuli and intact objects (Figure 1). The stimuli were presented in random order at four presentation frequencies (60Hz, 30Hz, 20Hz, 5Hz) to target different levels of visual processing (Grootswagers et al., 2019; Robinson et al., 2019). The stimuli were developed by Long et al., (2017), and obtained from https://osf.io/69pbd/ (Long et al., 2017, 2018). Continuous EEG was recorded during the streams and cut into overlapping epochs based on the onset of each stimulus within the streams. The epoched data were subjected to a multivariate decoding analysis, similar to previous work that decoded individual images in fast presentation streams (Grootswagers et al., 2019; Robinson et al., 2019).

To investigate how image representations differed between the texform and intact versions, we obtained cross-validated classifier performance between all pairwise texform images (Figure 2A), and all pairwise intact object images (Figure 2B). The differences between texform and intact object decoding accuracies (Figure 2C) showed evidence for no difference in the initial response (up to around 150ms), but higher accuracies for intact objects after that. These differences were localised in occipito-temporal areas (Figure 2D). Both texforms and intact objects were decodable from around 90 ms after stimulus onset at all presentation frequencies, characteristic of early stages of visual processing (Carlson et al., 2013; Cichy et al., 2014; Contini et al., 2017). Faster presentation frequencies resulted in lower peak decoding and shorter decoding durations, consistent with previous results showing that fast rates restrict visual processing (Robinson et al., 2019). In general, the image-level decoding results were similar between texforms and intact objects, apart from the intact objects at 5Hz, where a larger second peak was observed that was not apparent for the texforms. To further investigate the similarity between the underlying patterns, we performed cross-decoding where we trained on intact object patterns and tested on texform versions (Figure 2E), and vice versa (Figure 2F). The results of these analyses showed that the evoked patterns are sufficiently similar to allow for above-chance cross-decoding at all presentation frequencies.

**Figure 2.**
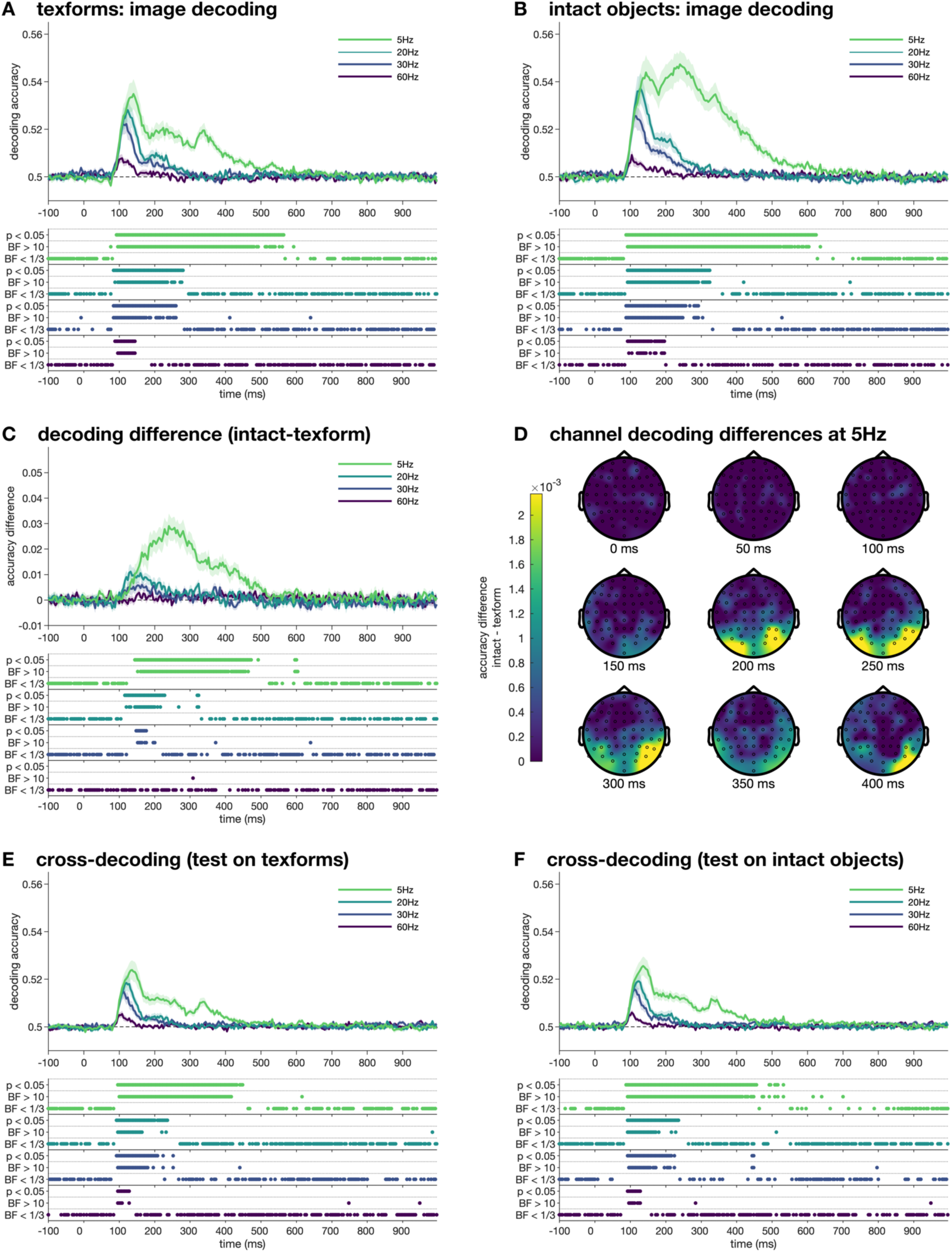
Decoding texforms and intact objects. **A.** Decoding between all texform image pairs. **B.** Decoding between all intact object image pairs. **C.** Difference between intact objects and texforms at each frequency. **D**. Channel searchlight maps at nine time points for the decoding differences at 5Hz. **E**. Decoding texforms, training the classifier on intact objects. **F.** Decoding intact objects, training on texforms. Different lines in each plot show decoding accuracy for different presentation frequencies over time relative to stimulus onset, with shaded areas showing standard error across subjects (N=20). Thresholded Bayes factors (BF) and p-values for above-chance decoding or non-zero differences are displayed below the plot for each frequency.

To investigate to what extent the visual features preserved in the texform versions of the objects drive categorical distinctions in activation patterns, we trained classifiers on decoding animacy from the stimuli, using an exemplar-by-sequence cross-validation approach to avoid overfitting to individual images (Carlson et al., 2013; Grootswagers et al., 2019). For texforms, above chance decoding of (featural) animacy was observed between 300ms and 400ms after stimulus onset (Figure 3A) for the 5Hz presentation frequency but was not evident for the faster presentation rates. Animacy decoding in intact object images, in contrast, was above chance for 5Hz, 20Hz and 30Hz (Figure 3B). Onset of animacy decoding was approximately 150ms for the 5Hz and 20Hz conditions, and at 220ms for the 30Hz frequency. The differences between texform and intact object animacy decoding accuracies (Figure 3C) highlight the substantial difference at 5Hz, which an exploratory searchlight suggested to be mainly located across occipito-temporal areas (Figure 3D). Cross-decoding (Figure 3E&F) showed that part of the animacy pattern generalised between intact objects and texforms. This result shows that the shared visual features between texforms and intact objects contributes to, but do not wholly explain, the categorical representation of animacy in the brain.

**Figure 3.**
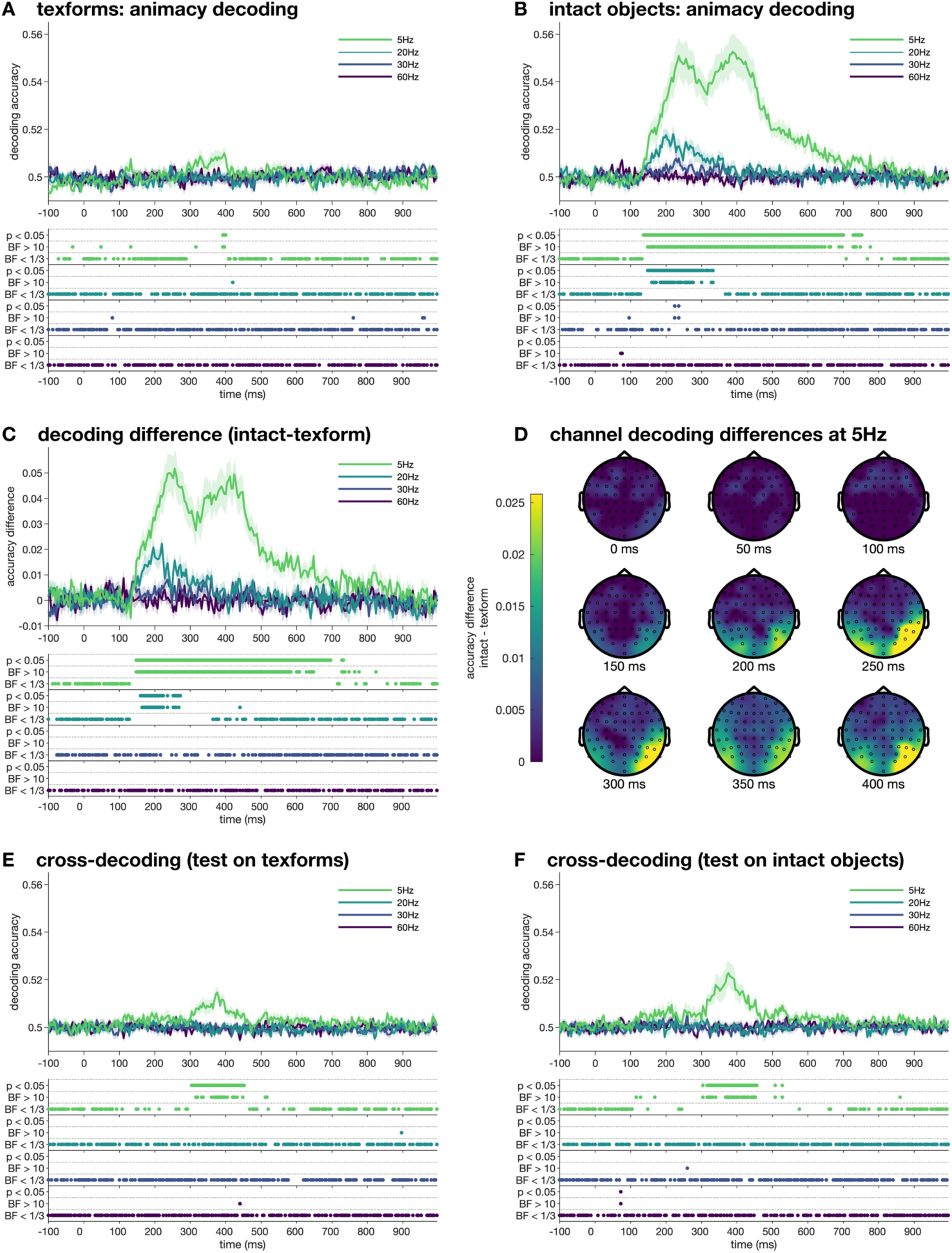
Decoding animacy from texforms and intact objects. **A.** Decoding the animacy of the texform images. **B.** Decoding the animacy of the intact objects. **C.** Difference between intact objects and texforms at each frequency. **D**. Channel searchlight maps at nine time points for the decoding differences at 5Hz. **E**. Decoding animacy from texforms, training the classifier on intact objects. **F.** Decoding animacy from intact objects, training on texforms. Different lines in each plot show decoding accuracy for different presentation frequencies over time relative to stimulus onset, with shaded areas showing standard error across subjects (N=20). Thresholded Bayes factors (BF) and p-values for above-chance decoding or non-zero differences are displayed below the plot for each frequency.

In the final analysis, we asked if the categorical representation of real-world size emerges similarly for intact and texform versions of objects. An exemplar-by-sequence cross-validation approach was used to decode real world size (small versus large objects) for the texform and intact objects. At none of the presentation rates was (featural) real-world size decodable from the texform stimuli (Figure 4A). Real world size of the intact object images was decodable for 5Hz and 20Hz frequencies (Figure 4B). The differences between texform and intact object size decoding accuracies are shown in Figure 4C. Cross-decoding showed evidence for no shared pattern of object size between intact objects and texforms (Figure 4E&F). Combined, the animacy and size decoding results show a fundamental difference in how conceptual categories emerge for intact objects and their scrambled counterparts.

**Figure 4.**
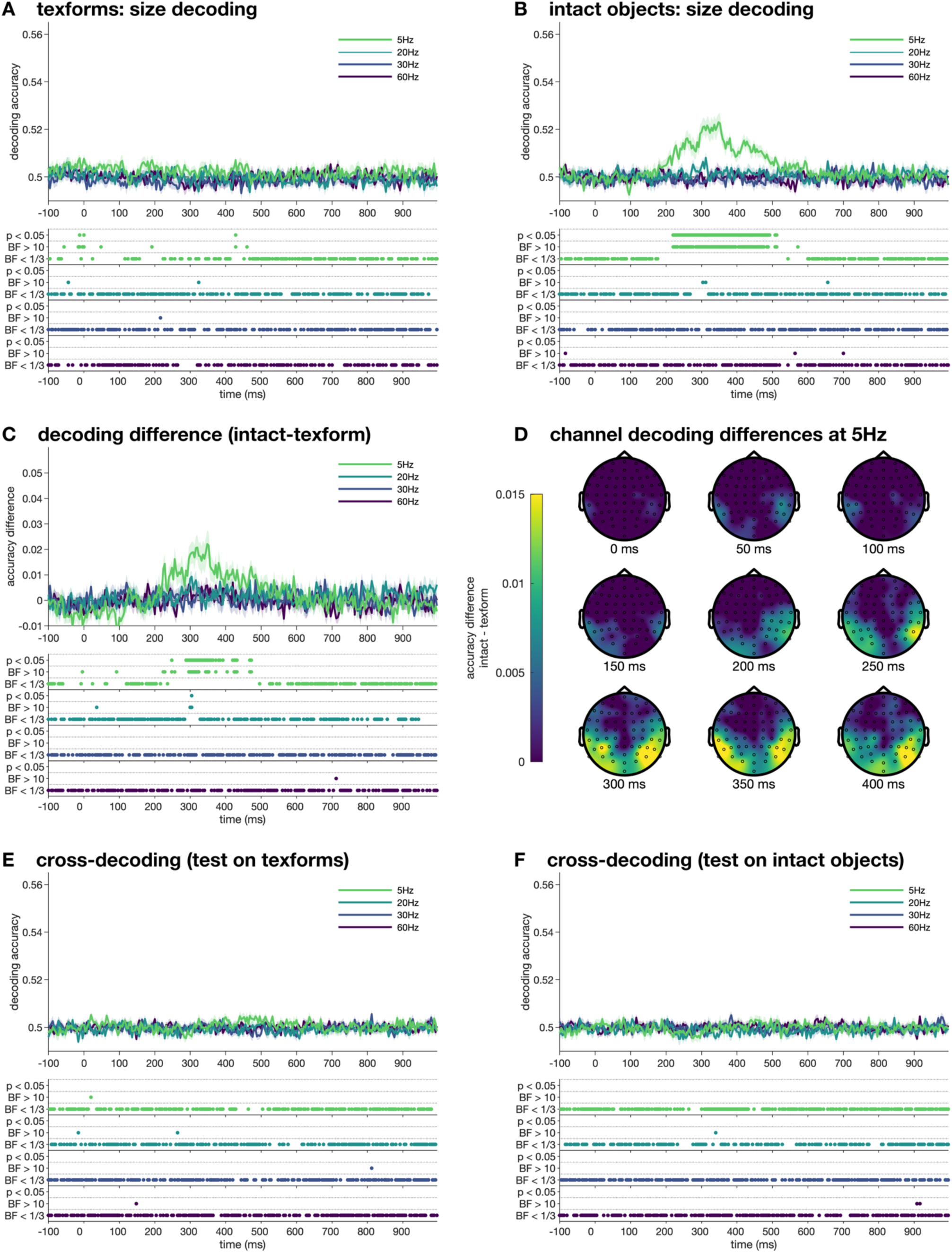
Size was only decodable from intact objects. **A.** Decoding the real-world size of the texform images. **B.** Decoding the real-world size of the intact objects. **C.** Difference between intact objects and texforms at each frequency. **D**. Channel searchlight maps at nine time points for the decoding differences at 5Hz. **E**. Decoding size from texforms, training the classifier on intact objects. **F.** Decoding size from intact objects, training on texforms. Different lines in each plot show decoding accuracy for different presentation frequencies over time relative to stimulus onset, with shaded areas showing standard error across subjects (N=20). Thresholded Bayes factors (BF) and p-values for above-chance decoding or non-zero differences are displayed below the plot for each frequency.

## Discussion

In this study, we assessed the contribution of mid-level features to high level categorical object representations using a combination of fast periodic visual processing streams and multivariate EEG decoding. We used images of intact and texform versions of objects from a previously published study (Long et al., 2018) and found that their neural representations were similarly distinct at the image level. In contrast, the decoding accuracies of the original categorical distinctions of animacy and real-world size varied markedly across the texform and intact versions of the objects. The patterns of neural activity evoked by animate and inanimate intact objects were decodable during a larger time period than their texform versions, suggesting the temporal dynamics of animacy-like patterns varied between intact and texform versions despite their shared mid-level visual features. In addition, the animacy of intact objects was decodable at 5Hz, 20Hz and 30Hz, but texforms were only decodable at 5Hz. Higher level categorical brain regions exhibit larger responses to slower presentation rates relative to faster rates (McKeeff et al., 2007), and we previously found that slower object presentations reached higher, more abstract levels of visual processing (Grootswagers et al., 2019; Robinson et al., 2019). Thus, the absence of animacy decoding for texform objects at faster presentation rates indicates that higher level processing was required for the animate/inanimate distinction in texform stimuli. Moreover, a clear double-peak structure was observed for decoding the intact objects at 5Hz, but not for the texforms. This could reflect an additional conceptual processing step that is unique to intact objects presented at a frequency that allows reaching conceptual processing stages. We interpret these findings as evidence that shared visual features between texforms and intact objects contribute to, but do not wholly explain, the categorical organisation of animacy in the brain.

Our results corroborate fMRI results using the same stimuli showing that texforms and intact objects generated similar categorical representations along the visual hierarchy but that the recognizable images generated stronger category responses (Long et al., 2018). The current results further show a clear difference in the temporal dynamics of animacy representations within the visual system for featural versus conceptual object representations. At faster presentation rates, animacy decoding was observed in intact objects but not in texforms, indicating that intact objects promote categorical representations with limited processing. It is important to note that the intact objects were shown only in the second half of the experiment, which might have contributed to better animacy decoding for intact objects. However, image-level results were similar between texforms and intact objects, which suggests that the experimental paradigm was not wholly responsible for the differences in categorical-level decoding. Together, these results suggest that brain responses to intact objects contain additional animacy category information over and above the statistical visual regularities present in the texforms.

The texform scrambling process was used to render images unrecognisable at the individual image level, while maintaining featural image statistics. Some low-level visual information may have been lost in the scrambling process, such as shape and curvature information, which is a strong cue for animacy (Levin, Takarae, Miner, & Keil, 2001; Schmidt, Hegele, & Fleming, 2017; Zachariou, Giacco, Ungerleider, & Yue, 2018). In MEG and EEG decoding studies, classification can be strongly driven by differences in object shape (Proklova et al., 2019), and silhouette similarity is often a strong predictor of the similarities between the earliest neural responses (Carlson et al., 2013; Grootswagers et al., 2019; Teichmann, Grootswagers, Carlson, & Rich, 2018; Wardle, Kriegeskorte, Grootswagers, Khaligh-Razavi, & Carlson, 2016). It is also important to note that while the texform images are not recognisable at the individual level, they can still be categorised (e.g., for animacy) above chance (Long et al., 2017). Human categorisation accuracies on these images was found to be predicted by the amount of curvilinear and rectilinear information in the image (Zachariou et al., 2018). Yet, even if intermediate visual features are sufficient to classify conceptual categories above-chance behaviourally, our results suggest that this is only possible given sufficient processing time.

These findings support the notion that large-scale categorical organisations in the visual system are to some extent driven by mid-level visual features. However, if concepts were decodable using only brain responses to mid-level feature, then this would predict above-chance decoding of concepts also at faster frequencies for the texforms. This was not the case in our results. Instead, we only observed animacy decoding for the slowest (5Hz) presentation frequency, which suggests that the conceptual animacy category only emerges from mid-level features after deeper processing. Thus, it could be the case that mid-level feature coding in early visual areas does not allow for concept decoding, but these features are “untangled” by higher visual areas into linearly separable categorical organisations (DiCarlo & Cox, 2007). This would mean that visual features could indeed drive the organisation in high level areas, but only given sufficient processing time for such untangling processes to complete. Furthermore, the speed of processing or information transfer to these higher visual areas could be modulated by the amount of evidence that supports the successful recognition of an object. For example, the intact objects have a well-defined outline that separates the object from the background, while the edges of the texforms are more blurred. This potentially could disrupt segmentation processes which in turn delays the amount of time it takes for information to reach higher level recognition stages. The results of this study therefore suggest a complex interplay between early and late stages of processing that ultimately manifests in more abstract categorical representations.

A limitation of the current study is that the animacy and size category boundaries of the stimulus set also define four subcategories, for example, most small objects are tools, and most big animals are mammals. Therefore, the results could be driven by these subcategories, rather than overall conceptual animacy and size organisations. Future work could explore this possibility, using stimulus sets that in addition match the subcategories within animacy and size categories. In addition, while it is important to disentangle perceptual features from conceptual representations, the two are inherently intermingled. Categorical organisations, such as animacy, are strongly represented partly because they share perceptual characteristics, which makes them easier to discriminate. Indeed, inanimate stimuli that share perceptual features with animate items evoke brain responses that are similar to other animate stimuli (Bracci et al., 2019). On the other hand, neural responses to animate stimuli that share characteristics with inanimate objects (e.g., a starfish) are more confusable with inanimate stimuli (Grootswagers, Ritchie, et al., 2017). Moreover, when stimuli are closely matched in shape, the activation patterns can become indistinguishable (Proklova et al., 2016, 2019). Together, these examples could be taken to suggest that the dichotomy of animacy should be revised to more closely reflect, for example, a continuous account of perceptual and conceptual animal typicality (Connolly et al., 2012; Contini, Goddard, Grootswagers, Williams, & Carlson, 2019; Grootswagers, Ritchie, et al., 2017; Iordan, Greene, Beck, & Fei-Fei, 2016; Sha et al., 2015; Thorat, Proklova, & Peelen, 2019).

In conclusion, we found that animacy was decodable from texform versions of objects, but that animacy of intact objects was more strongly decodable, and at faster presentation frequencies. Information contained in the texform versions of the objects thus not fully account for the distinct patterns of neural responses evoked by conceptual object categories. These findings suggest that complex interactions between lower and higher levels of visual processing mediate the representations of category, which has important implications for disentangling perceptual and conceptual representations in the human brain.

## Acknowledgements

This research was supported by an Australian Research Council Future Fellowship (FT120100816) and an Australian Research Council Discovery project (DP160101300) awarded to T.A.C. The authors acknowledge the University of Sydney HPC service for providing High Performance Computing resources. The authors declare no competing financial interests.

